# Locally induced traveling waves generate globally observable traveling waves

**DOI:** 10.1101/2025.01.07.630662

**Authors:** Kirsten Petras, Laetitia Grabot, Laura Dugué

**Affiliations:** Université Paris Cité, CNRS, Integrative Neuroscience and Cognition Center, F-75006 Paris, France; Laboratoire des systèmes perceptifs, Département d’études cognitives, École normale supérieure, PSL University, CNRS, 75005 Paris, France; Institut Universitaire de France (IUF), Paris, France

## Abstract

Cortical traveling waves have been proposed as a fundamental mechanism for neural communication and computation. Methodological uncertainties currently limit the interpretability of non-invasive, extracranial traveling wave data, sparking debates about their cortical origin. Studies using EEG or MEG typically report waves that cover large portions of the sensor array which are often interpreted as reflecting long range cortical waves. Meanwhile, invasive, intracranial recordings in humans and animals routinely find both local, mesoscopic waves and large scale, macroscopic waves in cortex. Whether the global sensor-array waves found with EEG/MEG necessarily correspond to macroscopic cortical waves or whether they are merely projections of local dynamics remains unclear. In this study, we made use of the well-established retinotopic organization of early visual cortex to generate traveling waves with known properties in human participants (N=19, m/f) via targeted visual stimulation, while simultaneously recording MEG and EEG. The inducer stimuli were designed to elicit waves whose traveling direction in mesoscopic retinotopic visual areas depends on stimulus direction, while leaving macroscopic activation patterns along the visual hierarchy largely unchanged. We observed that the preferred direction of traveling waves across the sensor array was influenced by that of the visual stimulus, but only at the stimulation frequency. Comparison between single-trial and trial-averaged responses further showed considerable temporal variation in traveling wave patterns across trials. Our results highlight that under tight experimental control, non-invasive, extracranial recordings can recover mesoscopic traveling wave activity, thus making them viable tools for the investigation of spatially constrained wave dynamics.

**Significant statement:** Non-invasively obtained time-resolved neuroimaging data is often thought to primarily reflect neural dynamics on the largest spatial scales. In the context of cortical traveling waves, this assumption can lead to a misinterpretation of spatio-temporal patterns observed in the sensor array. We here show that it is in principle possible that the global sensor array data is dominated by spatially constrained, local cortical traveling wave activity. Our findings crucially inform the ongoing discussion about the origin of traveling waves observed in surface recordings.

## Introduction

Brain activity recorded with electroencephalography (EEG) or magnetoencephalography (MEG) has repeatedly been found to form traveling waves across sensors. These waves, characterized by a monotonic phase shift in the direction of signal propagation across the sensors, are often thought to reflect cortical waves with equivalent temporal and spatial frequencies (Alamia et al., 2023; Alexander et al., 2006; Fakche et al., 2024). This view has led to speculations about the functional relevance of traveling waves, for example as the carrier of prediction errors in predictive coding frameworks (Alamia & VanRullen, 2019) or as determining perceptual rhythms across retinotopic space (Fakche & Dugué, 2024; Sokoliuk & VanRullen, 2016).

Recently, however, authors have questioned the interpretability of extracranially observed apparent traveling waves. They suggested that two spatially discrete dipole sources or multiple discrete clusters of time lagged activity could explain the observed data equally well (Orsher et al., 2023; Zhigalov & Jensen, 2023). Alternatively, local, mesoscopic traveling waves (covering short distances, up to a few cm) could project to the surface of the head and there be registered by the sensor array as seemingly global waves (Hindriks et al., 2014; Orczyk & Kajikawa, 2022).

MEG and EEG are non-invasive and offer full head coverage, making them popular tools for the study of traveling waves. Understanding the sources underlying the globally observed EEG/MEG activity is crucial for drawing meaningful inferences from the data. Source reconstruction of sensor array traveling waves as a means to differentiate between competing accounts is however precluded due to methodological limitations (see also Grabot et al., 2024): the spatial blurring and distortion of signals as they pass through diverse media before being recorded by distant sensors that is inherent to EEG/MEG data (van den Broek et al. 1998; Wolters et al. 2004) is here compounded by the fact that the most focal techniques available today rely on the use of spatial filters that necessarily obscure the spatial dynamics of traveling waves due to their inherent assumption of spatially stationary sources (Hillebrand & Barnes, 2005, 2011; Lin et al., 2008). Low signal-to-noise ratios in recordings and the resulting need for trial averaging might likewise obscure diffuse temporal dynamics and further complicate linking local to global traveling waves (Alexander et al., 2013). Alternatively proposed invasive intracranial recordings, such as electrocorticography, stereotactic EEG in humans or microelectrode arrays and voltage sensitive dye optical imaging in animals suffer less from the effect of volume conduction, but typically cover a smaller portion of the cortex. These techniques are suitable to record mesoscopic waves, but lack the dense and broad spatial coverage to represent macroscopic patterns traversing large distances, up to the size of the full cortex (but see Alexander and Dugué, (2024). Clarifying the spatial scales that non-invasive recording techniques with large spatial coverage can resolve in the study of traveling waves is therefore critical to addressing this significant gap in the literature.

Here, we make use of the retinotopic organization of early visual cortices to produce mesoscopic cortical traveling waves with known properties in localized cortical areas and assess their macroscopic effects in MEG-EEG recordings. Participants viewed a dynamic stimulus, adjusted for the cortical magnification of the primary visual cortex (V1) and designed to produce either a wave traveling in (periphery towards fovea), traveling out (fovea towards periphery) or a locally standing wave in retinotopic areas. Concurrent MEG-EEG was recorded and all participants underwent fMRI retinotopic mapping used to estimate wave propagation speeds in V1. We recovered propagation patterns from recorded data using optical flow analysis and relate these measures back to the known properties of the retinotopically-induced waves. With this, we assessed two competing hypotheses: (1) macroscopic traveling waves in the sensor array reflect mesoscopic traveling waves in cortex and (2) macroscopic traveling waves in the sensor array reflect sequential region activation (**Figure 1**). Our results are largely in line with the first hypothesis: the direction of the globally observed waves was dependent on the direction of the visually-induced wave, thus demonstrating that under tight experimental control, local waves *can* be picked up as a seemingly global wave in MEG-EEG sensors.

**Figure 1.**
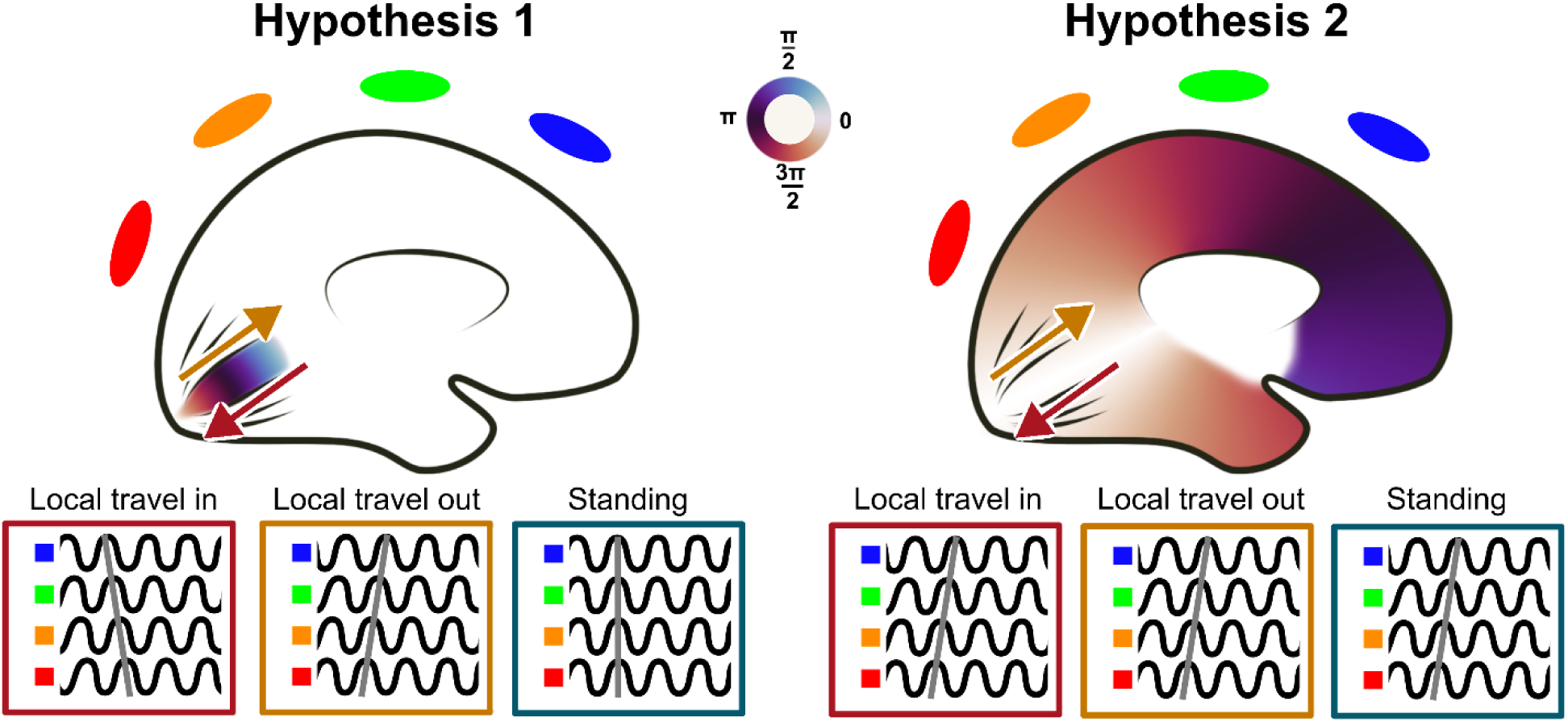
Two competing hypotheses on the cortical origin of globally observed traveling waves. Left, the locally induced mesoscopic wave projects to the surface of the cortex resulting in a global wave in the measurement array. Arrows indicate propagation direction of the induced mesoscopic wave. The brown-to-purple color spectrum indicates the phase distribution at an arbitrary point in time that underlies the global wave observed in the sensor array. Because of the larger distance traveled by the projection, the global wave might appear to be propagating considerably faster than the retinotopically constrained local wave(Hindriks et al., 2014). In this scenario, the direction of the stimulus (travel in vs. travel out) is expected to determine the direction of the globally induced wave. Subtle differences in individual cortical topography as well as sensor positions relative to the visual cortex can significantly impact observed wave directions across participants but should remain stable across trials within individuals. **Right,** the local stimulus triggers a sequential activation of (visual) cortical areas with set distance-dependent delays. The traveling direction of the induced mesoscopic wave (arrows) is the same as in the left panel, however, the phase map producing the observed global wave differs. Here, the oscillatory visual input results in oscillatory responses in each step of the visual hierarchy and the region-to-region delays result in a systematic phase shift between the oscillating populations. The stimulus direction is expected to be less impactful than the presence or absence of a stimulus altogether. An alternative explanation to both of those scenarios is that the globally observed wave results from a third mechanism such as global attentional processes and is not directly linked to the stimulus (see discussion).

## Materials and Methods

All data were originally collected for a different study and re-analyzed here. Full descriptions of stimuli, task and recording conditions can be found in the associated manuscript (Grabot et al., 2024). Here, we detail the relevant materials and methods for the present study.

### Participants

20 healthy volunteers (11 female, age M = 28 years, SD = 6) with normal or corrected to normal vision participated in the study. Participants were naive to the purpose of the study, provided written informed consent and were compensated for their participation. All data were recorded at the Paris Brain Institute (ICM), France. The protocol was in accordance with the Declaration of Helsinki (2013) and approved by the French Ethics Committee on Human Research (RoFOCAC, #ID RCB 2017-A02787-46). One participant was excluded a priori due to excessive noise in the MEG signal. A total of 19 participants’ data (10 female, age M = 28 years, SD = 5) entered analysis.

### Anatomical and functional MRI acquisition

Anatomical and functional MRI images were obtained using standard sequences on a 3T Siemens Prisma scanner. T1-weighted anatomical images, obtained with a FOV of 256mm and a voxel sixe of 0.8mm^3^ with no gap were used in conjunction with functional retinotopic mapping images at 2mm resolution to calculate, for each participant, the expected cortical propagation properties of the induced traveling wave in retinotopic primary visual cortex. Functional MRI data were preprocessed using fMRIPrep 20.2.1 version ((Esteban et al., 2019), RRID:SCR_016216) and the population receptive field model was produced using analyzePRF, a Matlab toolbox based on a compressive spatial summation model (Dumoulin & Wandell, 2008; Kay et al., 2013).

### Stimuli

All stimuli were created and presented using Matlab R2015B and Psychtoolbox 3.0.15(Kleiner et al., 2007). Stimuli were viewed on a gray screen with 1920*1080 pixels (79.5*44.5 cm; 32.5 x 18.3 °VA) at a viewing distance of 78 cm using a PROPixx projector at 120 Hz refresh rate.

Here, the data of three stimulus conditions were analyzed: traveling out stimulus (traveling from the fovea to the periphery), traveling in stimulus (same stimulus traveling from the periphery to the fovea) and standing stimulus.

The traveling stimuli were designed to elicit a traveling wave across the retinotopic space. The luminance of the screen was modulated according to equation (Eq 1), with a spatial frequency Fs = 0.05 cycles/mm of cortex, a temporal frequency Ft = 5 Hz and an initial phase shift. The wave amplitude and the offset (A = 0.5, c = 0.5, respectively) were chosen so that the luminance fluctuated between 0 (white screen) and 1 (black).

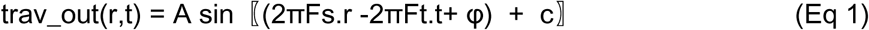

in which r corresponds to the radial distance from the center of the screen of a given pixel on the screen, after correction for cortical magnification.

The magnification inverse M^-1^ scaled with the eccentricity e of a given point on the screen (°VA, degrees of visual angle) and followed equation (Eq 2) with M0 = 23.07 mm/°VA, E2 = 0.75° VA (from Figure 9 in Strasburger et al., (2011), parameters from Horton & Hoyt, (1991)).

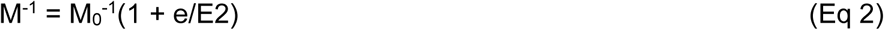

The temporal frequency of the traveling stimuli was chosen based on a previous study on traveling waves in the theta frequency range (Zhang et al., 2018). The spatial speed (0.05 cycles/mm at 5 Hz corresponds to a wave propagating at 0.1 m/s) also matched the empirically observed velocity of mesoscopic traveling waves (0.1-0.8 m/s, from Muller et al., (Muller et al., 2018); Fakche and Dugué, (Fakche & Dugué, 2024)).

For the standing stimulus, the luminance of the screen was modulated using equation (Eq 3) after adjusting for cortical magnification, with the same parameters as the traveling stimuli.

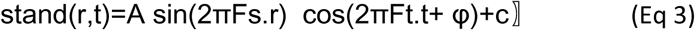

The traveling in condition followed Eq 4, with the same parameters as Eq 1.

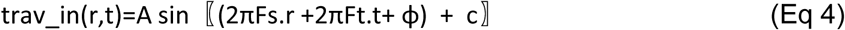

To create each stimulus, the luminance modulation described by the previous equation was then multiplied with a static carrier. The carrier corresponded to Gaussian white noise with a spatial frequency tuned to V1’s preferred spatial frequency, following equation (Eq 5):

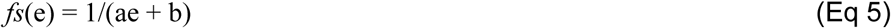

with *fs* the preferred spatial frequency (cycles/mm) in V1 for a given eccentricity e (°VA), a = 0.12, and b = 0.35 (values from Broderick et al., (Broderick et al., 2022)).

Binarization was then applied to maximize contrast. To avoid any adaptation effect to the static carrier, 10 different randomly created carriers were presented (one per block) in a pseudo-random order across participants. Within a block, the given carrier was presented in half of the trials in its original orientation and in the other half, rotated by 180°, in a pseudo-random order across conditions.

The final stimuli were obtained by multiplying the luminance modulator with the carrier, after rescaling between 0 (black) and 1 (white) as follows:

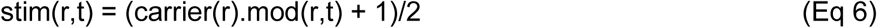

with mod being either *trav_out*, *trav_in* or *stand* and r the radial distance from the center of the screen of a given pixel.

### Experimental procedure

Participants fixated a red fixation dot at the center of the screen (0.05 °VA of diameter; Figure **2A**). Each trial consisted of a visual stimulus presented for 2 seconds, followed by a 2-second fixation interval. Participants completed 10 blocks (∼7min each) of 8 runs each. During each run, 8 trials were presented and each stimulus condition was repeated twice, in random order, resulting in 160 repetitions for each stimulus condition. To ensure participants attention to the stimulus, they performed a simple color change detection task at fixation (dot changing from red to yellow with a probability of 25%). Participants responded by pressing a key with their right hand after the run ended. The color change occurred in one out of 8 trials, anywhere between 0.5 and 1.8 seconds from stimulus onset and lasted 150ms.

**Figure 2.**
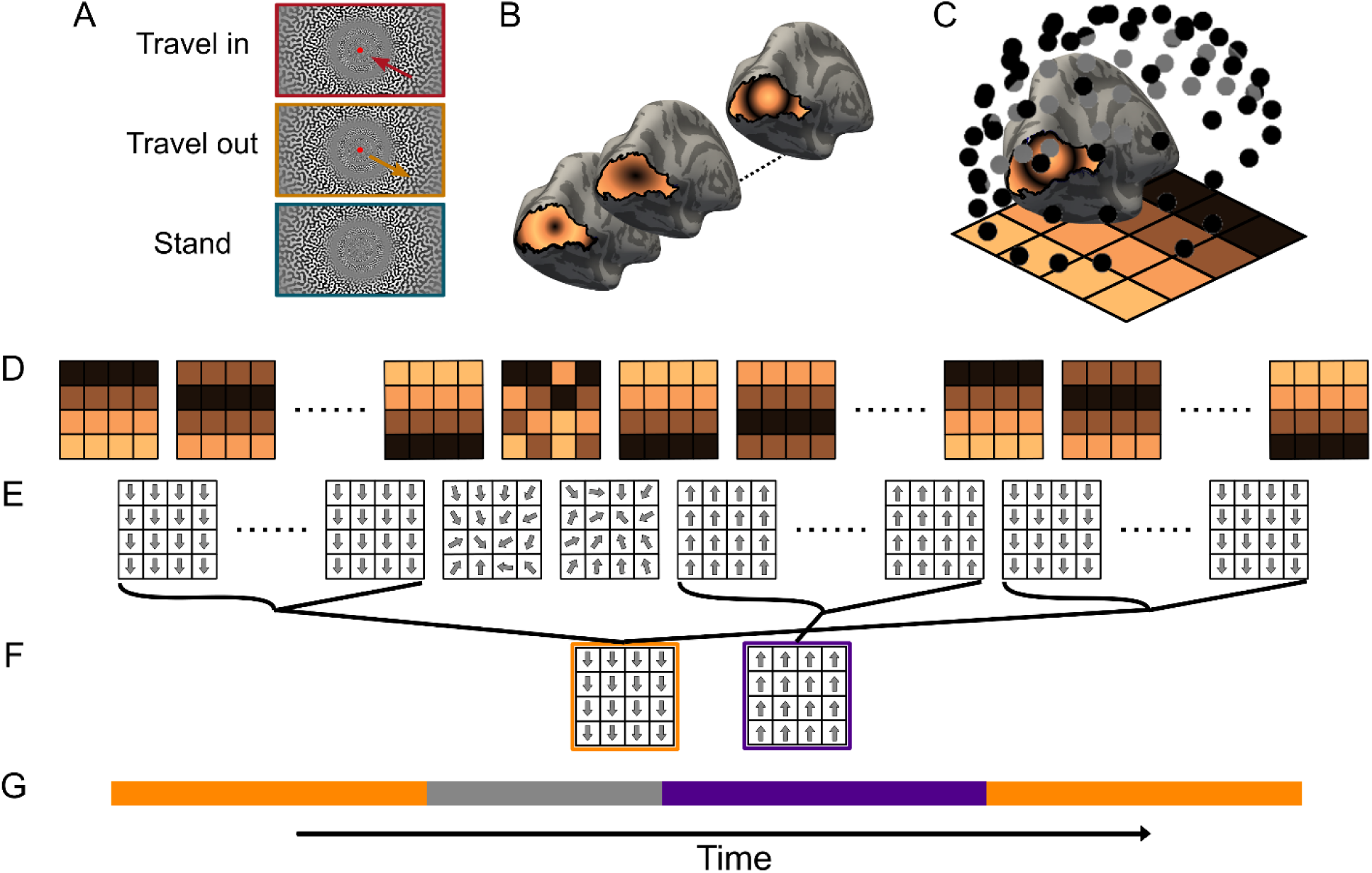
Motif extraction pipeline. **A.** Participants viewed a stimulus designed to produce a smooth traveling wave in retinotopic cortex (**B**). **C**. Concurrent EEG and MEG were recorded and sensor data were projected to a 2-dimensional surface and interpolated to a regular grid. **D.** Hilbert-phase timeseries were subjected to optical flow analysis. **E.** Temporally stable sequences of optical flow (flow vectors of sufficient magnitude pointing in the same direction for a minimum of one cycle of the frequency of interest) were extracted and **F.** averaged into re-occurring motifs. **G.** This procedure resulted in motif-coded timeseries of each individual trial. Here, orange indexes timepoints that displayed a motif with flow vectors pointing down whereas purple indexes flow vectors pointing up. Gray indexes timepoints where no spatio-temporally stable motif was identified.

Participants’ behavioral performance was assessed with dprime, computed by subtracting the z-transformed false alarm rate (FA) from the z-transformed hit rate (H). The average dprime was 1.26 ± 0.31 (H = 52 ± 5%, FA = 21 ± 5 %), indicating that participants were performing the task well above chance and successfully fixated the center dot.

### MEG-EEG recording

Electromagnetic brain activity was simultaneously recorded with EEG and MEG in a magnetically shielded room. MEG was collected using a whole-head Elekta Neuromag TRIUX MEG system (Neuromag Elekta LTD, Helsinki) in upright position. The system is equipped with 102 triple sensor elements (one magnetometer and two orthogonal planar gradiometers). EEG was recorded using an Easycap EEG cap compatible with MEG recordings (BrainCap-MEG, 74 electrodes, 10-10 system).

The MEG-EEG signal was recorded at 1 kHz sampling frequency with a low-pass filter of 330 Hz. A high-pass filter at 0.01 Hz was applied. Horizontal and vertical electro-oculogram (EOG) were collected using two electrodes near the lateral canthi of both eyes and two electrodes above and below the dominant eye. Electrocardiogram (ECG) was recorded with two electrodes placed on the right collarbone and lower left side of the abdomen. The ground electrode for the EEG was placed on the left scapula and the signal was referenced to connected electrodes on the left and right mastoids.

Participants’ head position was measured before each block using four head position coils (HPI) placed on the EEG cap over the frontal and parietal areas.

### Simulations

To validate and optimize all analysis steps and choices prior to the analysis of recorded data, we produced a set of simulated trials and subjected them to the planned analysis pipeline. Simulations were initially generated on a 42 x 42 regular grid. We generated 2 basic trial types (wave initially traveling top to bottom of the grid and wave traveling bottom to top), with 10 variations of gaussian random noise at two SNRs each, resulting in 20 trials in total. The trials were constructed as a 1000ms period of 1/f noise, followed by a plane wave traveling in either top-to-bottom or bottom-to-top direction for a duration of 1000ms. After 2 seconds of total simulation time, another noise interval of 100 ms followed, after which the wave changed direction (opposite to the original direction) and continued on for another 900ms. Plane waves were created as sinusoidal signals propagating across the two-dimensional grid where the signal at each grid point (x,y) and time t is described by the equation:

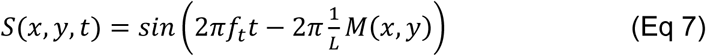

where 𝑓𝑡 is the temporal frequency and 𝐿 = 1/𝑓𝑠 is the wavelength, with 𝑓𝑠 being the spatial frequency of the wave (in cycles per grid-size). 𝑀(𝑥, 𝑦) is the spatial gradient function representing the phase shift in the direction given by the wave’s orientation (determined by the angle θ between the wave’s propagation direction and the positive x-axis). M(x,y) is calculated by applying a rotation matrix R to the grid coordinates (x,y). Overall, the plane waves generated in this manner exhibit sinusoidal oscillations modulated by the spatial gradient M(x,y), that represents the phase shift across the grid produced by the wave’s propagation. Generated waves had a temporal frequency of 𝑓𝑡 = 5 Hz, a spatial frequency 𝑓𝑠 = 0.8 cycles per grid and were sampled at 250Hz. In addition, we added scattered local oscillators at ∼20% of grid points for half of the generated trials. These were individual grid points, oscillating at 10 Hz with either random phases relative to each other (50% of trials) or globally coherent phase (50% of trials), that were mixed with the original waves at an SNR of 1. The affected grid points were thus intrinsically oscillating at an alpha frequency but could also be modulated by the passing wave. Finally, gaussian random noise at one of two SNRs (0.8; 1.8) was added to all trials.

To include a basic test of the interpolation of 3D sensor positions to a 2D regular grid that is necessary in real data sampled from irregular sensor positions, we replicated the original x,y - grid data, stacked it along a new z-dimension and scaled it to encompass a standard 74 channel EEG layout. We then sampled from this 3-dimensional array at the locations of the sensor positions, assuming a spatial extent of each sensor of 5mm (diameter). This resulted in a data array with the same shape as could be expected from recorded data and formed the basis of all further processing. All subsequent processing was identical between Simulations, EEG data and MEG data.

### MEG-EEG pre-processing

Noisy MEG sensors were excluded after visual inspection of raw data. Signal space separation (SSS) was then applied to the raw continuous data to decrease the impact of external noise (Taulu & Kajola, 2005). The head position measured at the beginning of each block was realigned relative to the block coordinates closest to the average head position across blocks. SSS correction, head movement compensation, and noisy MEG sensor interpolation were applied using the MaxFilter Software (Elekta Neuromag). Using the MNE-Python suite (Gramfort et al., 2014), noisy EEG sensors were interpolated using spherical spline interpolation. Ocular and cardiac MEG-EEG artifacts were reduced using ICA decomposition. MEG-EEG data were aligned to the detected ECG and EOG peaks. The ICA components capturing ocular and cardiac artifacts in brain responses were computed on these time-aligned signals separately for EEG and MEG recordings. The correction consisted in rejecting ICA components of brain data most correlated with the ECG and EOG signals. The rejection criterion was defined by a Pearson correlation score between MEG-EEG data and the ECG- and EOG-signals with an adaptive z-scoring (threshold = 3). All outcomes were verified by visual inspection. The EEG signal was re-referenced to the common average.

Data were low-pass filtered using a windowed zero-phase shift finite impulse response time-domain filter (53 dB stopband attenuation, 20 Hz transition bandwidth, passband edge 80 Hz, cutoff frequency 90 Hz, filter length .165s) to avoid aliasing in the subsequent decimation. Data were then epoched into segments spanning -1 to 2 seconds around stimulus onset and decimated to a final sampling rate of 250 Hz following best practices (https://mne.tools/stable/auto_tutorials/preprocessing/30_filtering_resampling.html#best-practices).

All further analyses were carried out using the Wave Space software (see **Code and data availability** section).

### Frequency decomposition

To isolate oscillatory data components, we used a narrowband filter, followed by a Hilbert transform. Frequencies of interest (FOI) were 5 Hz (frequency of the stimuli), and each participant’s individual alpha frequency. To determine individual alpha frequencies, we extracted the spectral peak in the alpha band (8-12 Hz) from the pre-stimulus period while controlling for the aperiodic component (the typical 1/f slope) of the power spectrum using the FOOOF toolbox (Donoghue et al., 2020). For simulations and cases where no clear spectral peak was found (in 2 participants’ data), a frequency of 10 Hz was used. Data segments were then band-pass filtered with cutoffs set to FOI -1 > FOI < FOI +1 (hamming windowed non-causal finite impulse response filter with 0.0194 passband ripple and 53 dB stopband attenuation. Lower transition bandwidths were 2 Hz (5 Hz center frequency of passband) and ∼2.22 Hz (individual alpha peak center frequency of passband). Upper transition bandwidth were 2 Hz and ∼2.72 Hz, respectively. Filter length were 1.652 s for theta and ∼1.492 s for alpha. Subsequently, the Hilbert transform was computed after spatial interpolation (see next step). Non-causal FIR filters provide precise frequency band control and preserve phase, which is well-suited to extracting narrowband oscillatory components, but can be prone to ringing and edge artifacts (Widmann et al., 2015). Subsequent phase analysis was focused on the steady-state responses to visual flicker stimulation, excluding the initial stimulus onset and trial edges.

### Spatial interpolation

The resulting narrowband filtered time-series of all four data types (EEG, MEG magnetometers, MEG gradiometers and simulations) were interpolated to a 2D regular grid as follows: Geodesic distances between digitized 3D channel locations along the curved surface they form were projected onto a 2D plane using multidimensional scaling (MDS), a well-established dimensionality reduction technique that respects the spatial relations between components (Borg & Groenen, 2005). In the neuroscience literature, MDS is most commonly used to express similarity and dissimilarity across different measurement modalities (Kriegeskorte et al., 2008), but has also previously been used to determine 2D relative electrode positions for cochlear implants (Vanpoucke et al., 2011). Here, the MDS-derived relative 2D positions were rotated to match the original orientation of the sensor array and interpolated onto a regular grid spanning the spatial extent of the MDS solution. We used spline interpolation, tiling the space between the most posterior and the most anterior electrode in 18 steps. This procedure ensured a smooth and continuous representation of the sensor data, with the same number and arrangement of spatial bins for EEG and MEG data. Gradiometer data was treated separately for the two sets of planar gradiometers.

### Optical flow analysis

To characterize consistent shifts in the oscillatory phase of electrical potentials across the space of the sensor array (i.e., traveling waves), we used a simplified version of the Horn-Schunk algorithm for dense optical flow estimation (Horn & Schunck, 1981). The original Horn-Schunk approach is based on globally minimizing both the brightness constancy assumption (i.e., brightness remains constant between consecutive frames) and the smoothness assumption (i.e., neighboring pixels have similar motion). Those can be written as

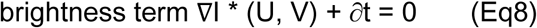

where ∇I represents the gradient of the image intensity at a pixel location, U and V are the horizontal and vertical components of the optical flow at that pixel and ∂t is the temporal derivative of the image intensity at that pixel,

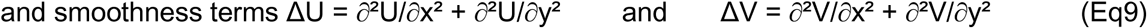

where ΔU and ΔV are the Laplacians of U and V, respectively and ∂x and ∂y are the partial derivatives with respect to the x and y spatial coordinates.

Deriving the optical flow can then be implemented by minimizing the objective function

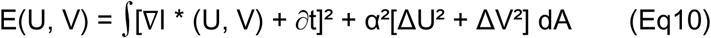

through the iterative updating of the flow field (U,V).

Here, ∫ represents the spatial integral over the entire image area, α is a regularization constant that controls the trade-off between brightness constancy and smoothness terms and dA represents the differential area element. The update step in each iteration is derived by solving the Euler-Lagrange equations for E(U, V). The final flow field (U, V), obtained after convergence of the optimization process, represents the estimated optical flow for each pixel location in the image. For computational efficiency, the updating step approximates the objective function’s minimization by iteratively updating the flow field based on local derivatives and averages rather than globally integrating over the entire image domain. The regularization parameter α determines the spatial smoothness of the resulting flow fields with smaller values producing finer-grained, but potentially noisy vector velocity patterns, whereas too large values can over-smooth the data and potentially miss local dynamics. Townsend and Gong ((2018)) recommend values in the range of 0.1-20, dependent on the spatial sampling frequency and the levels of noise in the data. Our data from both MEG and EEG recordings was interpolated to a grid representing ∼17mm inter-contact distances. We therefore chose α to equal 0.1. Optical flow analysis was applied separately to the phase maps of EEG and the two types of MEG data, with the two sets of gradiometers calculated independently and their optical flow maps combined as the sum of the individual flow vectors.

While the flickering stimulus presented to participants was expected to induce a strongly phase-locked response at the flicker frequency (5 Hz), previous research has shown that trial-by-trial variability in both phase and amplitude measurements is substantial, especially for spontaneous oscillations (here: alpha) (Stein, Gossen, and Jones 2005), and can be linked to behavior (Busch et al., 2009; Dugué et al., 2011; Fakche et al., 2022; Mathewson et al., 2009). Thus, to preserve potentially relevant trial-by-trial dynamics, we calculated the optical flow for each trial separately. In order to compare single-trial derived optical flow patterns to more commonly applied trial averaged analyses, we repeated the optical flow calculation on trial averaged phase estimates by using the circular average of each grid point over time.

### Flow field motifs

To summarize the resulting high-dimensional flow fields (frequency band x participant x trial x 2 spatial dimensions x time) into recurrent, temporally stable patterns (frequency band x participant x trial) - hereafter referred to as “motifs,” we followed a sequence of algorithmic steps:

For each trial, we calculated the cosine similarity, defined as the normalized scalar product between flow vectors at corresponding grid points but subsequent time points within a sliding time-window of minimally one cycle of the frequency of interest (200 ms for the 5 Hz component and ∼100 ms for alpha depending on the exact frequency for each given participant). A temporally stable sequence was initiated with a single template frame (flow field at a single time point). Additional frames were added to this sequence if they met the following conditions: (1) The cosine similarity between the current frame and the template frame (the first frame in the sequence) exceeded a threshold (.85, corresponding to ∼31.8°) in at least 40% of all grid points. (2) The flow vectors at positions where the cosine similarity exceeded threshold were of sufficient magnitude (> .1).

Any frame failing to meet the conditions for the current sequence, became the template frame for a new sequence. This process was repeated until all frames in the trial were processed. After all temporally stable sequences within a trial had been identified, sequences were compared to each other, within and across trials and participants (merge threshold = 0.8, corresponding to ∼36.9°), in a similar fashion to detect repeating motifs.

The exact parameters were determined from simulated data. Applying thresholds for oscillatory power/phase-locking did not qualitatively affect the final outcome, so for simplicity, only non-thresholded data is shown. The accompanying code contains options for both.

### Statistical analysis

We statistically tested the influence of our independent variables (conditions) on motif prevalence using chi squared statistics with two factors (condition, with 3 levels *travel in*, *travel out* and *stand* and 2 time-periods *pre-stim* and *stim*, see **Figure 5**). To avoid the initial stimulus onset response, the pre-stim and stim periods were defined as -0.5 to -0.01 s and +0.5 to 1.490s, respectively. The 6 most common motifs (across all conditions collapsed) entered the analysis. Separately for each sensor type, we counted motif occurrences across trials for each time point, participant and condition. These counts were averaged across time points separately for the two time periods.

To further assess whether a condition was associated with specific motifs, two additional analyses were carried out on motif counts. First, a linear mixed model was fitted, with condition (*travel in*, *travel out* and *stand*) and period (*pre-stimulus onset*, *post-stimulus onset*) as predictors, and participants as random effect. For simplicity, we restricted this analysis to the two most common motifs per modality (as they account for 78%, 77%, and 48% of the motif counts across time points, for EEG, gradiometers, and magnetometers, respectively).

To investigate which motifs, for a given participant, discriminate between traveling in and out conditions, we considered each motif’s count across the time points of the stimulation period, for each trial, condition and participant. These counts were normalized by the number of time points during stimulation to obtain a ratio. For each participant, we then performed a Lasso logistic regression on condition (travel in, travel out) with the six most frequent motifs as predictors. The L1 regularizer, chosen to favor weights sparsity, was estimated using 10-fold cross-validation on a training set corresponding to 60% of the data. It was taken as the value minimizing cross-validated errors. The classification accuracy was calculated on the testing set (remaining 40% of the data). All statistical analyses were carried out using R, including the glmnet package for the Lasso regression (Friedman et al., 2010).

Polar histograms of the direction of the preferred motif for each participant and condition were created as the average over all time points, across trials, matching the motif.

### Wave velocities

The velocity of waves corresponding to each motif was estimated by projecting optical flow vectors onto the primary direction of wave propagation for the respective motif. The magnitudes of the resultant vectors were then averaged over time and space, and adjusted using the geodesic distance along the sensor array and the data sampling frequency. This process provided an estimate of wave velocities specific to each motif.

Magnitudes of optical flow vectors are often used as a proxy for velocity (Gutzen et al., 2024). Note, however, that the smoothness parameter directly influences the flow vector magnitudes. Higher values of α, as well as large displacements, can lead to severe underestimation of velocities. Conversely, in regions with little or no motion, noise can cause an overestimation of velocity, thus requiring sufficient smoothing of noisy data (see Barron et al. (1994) for detailed performance measures). Additionally, the spatial interpolation of 3D sensor positions to a regular 2D grid necessarily introduces spatial distortions that affect velocity estimates from phase lags. We confirmed this by estimating velocities of the simulated waves, for which ground truth in the original array was known. Indeed, the optical flow overestimated velocity (true velocity: 1.67 m/s, velocity from vector magnitude ∼3.2 m/s). Nevertheless, the relative estimated velocities across conditions (estimated with the same optical flow parameters under the same spatial interpolation conditions), can provide some indication about the similarity between the travel paths. A global projection of the same velocity mesoscopic waves would, under similar noise conditions, result in very similar velocities for waves traveling in one or the opposite direction. Conversely, region sequencing should produce similar velocities for waves of all conditions traveling up the region hierarchy, but not those traveling down the hierarchy. Those are expected to follow a different propagation path, with different inter-region delays (Bullier, 2001; Felleman & Van, 1991; Markov et al., 2014; Van Essen & Maunsell, 1983). For comparison, we estimated the induced wave’s propagation velocity in primary visual cortex. For each visual angle, we calculated the ratio between the phase shift of the two furthest apart V1 voxels (i.e., at the two extremities of the eccentricity range) and the cortical Euclidian distance between these voxels. The individual spatial frequency was calculated as the average across all visual angles and across both hemispheres and velocity was estimated as the temporal frequency of the induced oscillation divided by the spatial frequency.

### Raw trial averaging versus motif averaging

In typical trial averaging, timeseries are averaged over multiple trials of the same condition while matching timepoints with respect to some trial event (here, stimulus onset). However, response components with inconsistent properties across trials are often not well represented by trial averaging and high amplitude components typically dominate the resulting signal (Mouraux & Iannetti, 2008). Decomposing data into matching components, or averaging over data segments that are more likely to contain stereotypical responses + gaussian white noise can mitigate this problem. Here, to highlight the effect of trial averaging of spatio-temporally rich data, we compared motif counts between averaged and non-averaged data. To create motif timeseries from trial averaged data, we averaged the raw signals of individual trials before narrowband filtering and extracting the analytic signal, separately for participant and stimulus condition, and subjected the resulting phase-timeseries to optical flow analysis. The following motif-extraction pipeline was the same as for single trials. Note that the motifs extracted from averaged data, as well as their order (sorted by their frequency of occurrence) do not necessarily match those from single trials. We then compared, for each participant, each timepoint of each individual trial’s UV map to the averaged motif timeseries of that same participant. For this, each motif extracted from the averaged data was treated as a template that individual timepoints’ UV maps were compared to as described above.

Similarly, average motif velocities were calculated as the average magnitude of the UV vectors projected onto the principal axis of the motif (main travel direction), estimated separately for all individual time points displaying the motif and compared with the velocities of the non trial-averaged data.

### Code and data accessibility

All analyses were carried out using the MNE python package (Gramfort et al. 2014, available under https://mne.tools/stable/index.html), the Wave Space python package (Petras et al., in prep., alpha version provided with accompanying code, the finalized package will be available on github upon publication of the corresponding manuscript) and custom code, available under https://github.com/kpetras/LocalGlobalWave. Anonymized data will be openly available on Zenodo upon publication of the original study (Grabot et al. 2024).

## Results

We investigated the global spatiotemporal patterns, measured with concurrent EEG and MEG recordings, induced by visually-driven, mesoscopic oscillatory traveling waves. We set out to differentiate between two accounts of the signal underlying the observed macroscopic waves in sensor arrays: 1) global projection of mesoscopic waves and 2) sequential activation of brain regions (**Figure 1**).

First, we found temporally sustained patterns of optical flow in all individual participants, in most trials (**Figure 3, Suppl. Figures 1&2** for individual participants’ results). We then grouped optical flow maps into temporally stable motifs. At the stimulation frequency (5 Hz), we found a difference in anterior-to-posterior vs. posterior-to-anterior motif prevalence between visual traveling wave directions (visual stimulus traveling toward vs. away from the fovea) in EEG data. Traveling toward the fovea (*travel in*) was associated with a higher frequency of anterior-to-posterior motifs and traveling away from the fovea (*travel out*) with a higher frequency of posterior-to-anterior motifs (**Figure 3, top row**; 60% more anterior-to-posterior during the travel in stimulation period). In gradiometer data, a similar effect of stimulation conditions was observed, albeit with travel directions rotated by 90 degrees, i.e., travel in was associated with 65% higher counts of right-to-left flow patterns and travel out with higher counts of left-to-right flow patterns (**Figure 3, middle row**). In magnetometer motif counts, condition differences showed the same trends as in gradiometer data, with 54% higher counts for right-to-left motifs (**Figure 3, bottom row**). The retinotopic standing wave had no significant effect on motif counts in any of the modalities. Statistical testing used a linear mixed model on the ratio motif A/motif B (the two most frequent motifs, averaged over all participants and conditions) which showed a main effect of condition for magnetometer and EEG (magnetometer: F = 3.94, p = 0.030; gradiometer: F = 1.85, p = 0.172; EEG: F = 9.36, p = 5.e-4), a main effect of period for magnetometer and gradiometer (magnetometer: F = 4.56, p = 0.049; gradiometer: F = 9.96, p = 0.005; EEG: F = 0.08, p = 0.784) and a significant interaction for magnetometer and EEG (magnetometer: F = 4.32, p = 0.022; gradiometer: F = 1.59, p = 0.218; EEG: F = 13.74, p = 4.1e-5).

**Figure 3.**
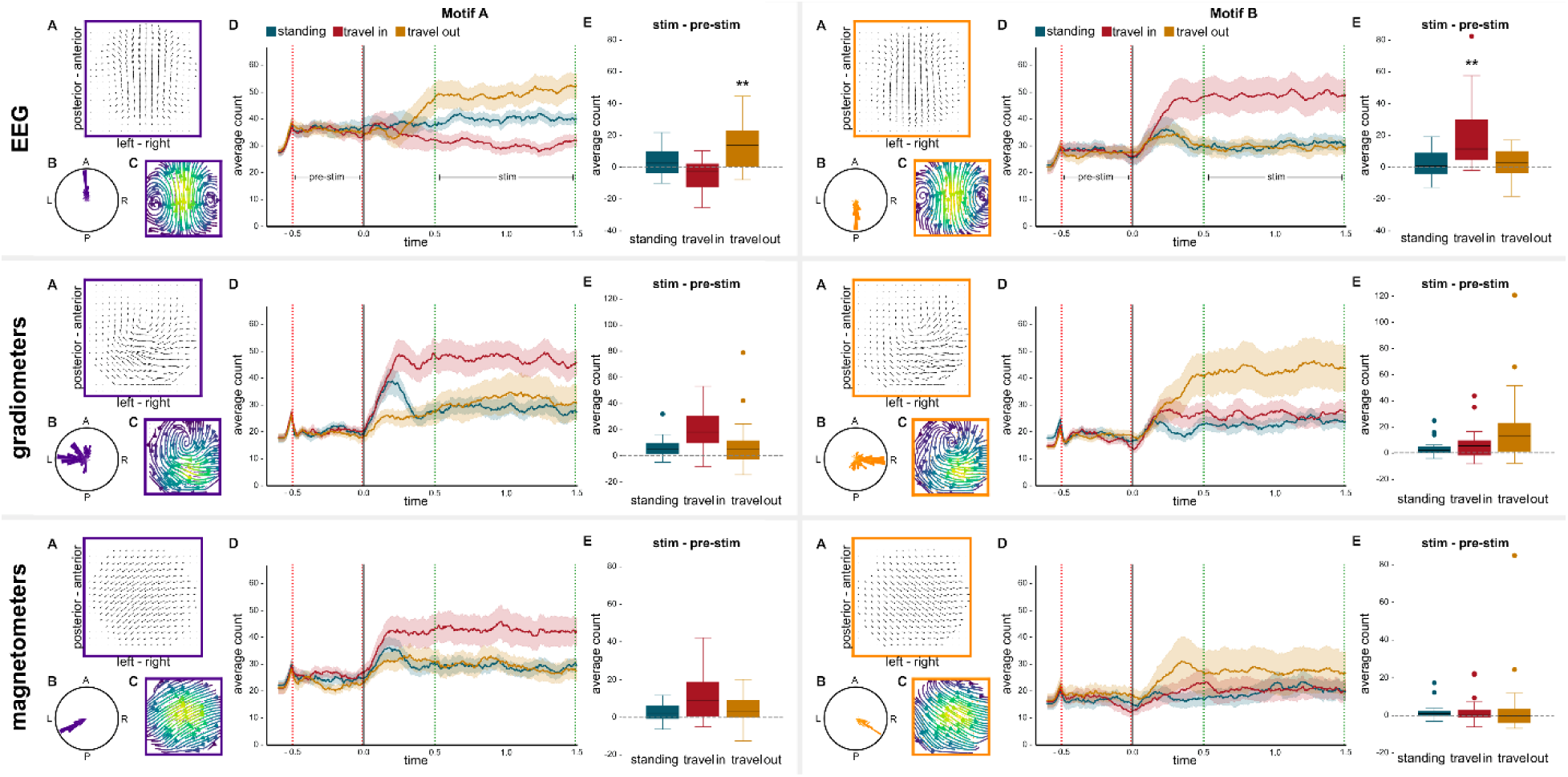
Average counts per condition for the two most frequent motifs (Motif A and Motif B) at 5Hz. The three rows show EEG, MEG gradiometer and MEG magnetometer data, respectively. Each column corresponds to a motif. For brevity, only the two most common motifs per imaging modality are shown. **A.** Motif template. Arrows show the average direction of optical flow vectors over timepoints matching the template. Arrow length is scaled by temporal consistency. **B.** Polar histogram of A. **C.** Streamline plot of average optical flow pattern for the current motif. Hot colors correspond to higher consistency of flow vectors across (non-consecutive) time points. **D.** Average frequency of motif occurrence over time. Shaded error margins correspond to standard error of the mean. **E.** Box plots of average counts in stimulus - pre-stimulus period. Colored dots indicate outlier data.

Optical flow maps created from alpha frequency data showed no such condition differences in any of the sensor arrays (GRAD, condition: F=0.426, p = 0.656, period: F = 4.037, p = 0.060, interaction, F = 0.97, p = 0.389; MAG, condition: F=1.191, p = 0.316, period: F = 0.08, p = 0.93, interaction: F = 2.759, p = 0.078; EEG, condition: F=0.404, p = 0.671, period: F = 2.436, p = 0.139, interaction: F = 2.766, p = 0.079; see **Figure 4**).

**Figure 4.**
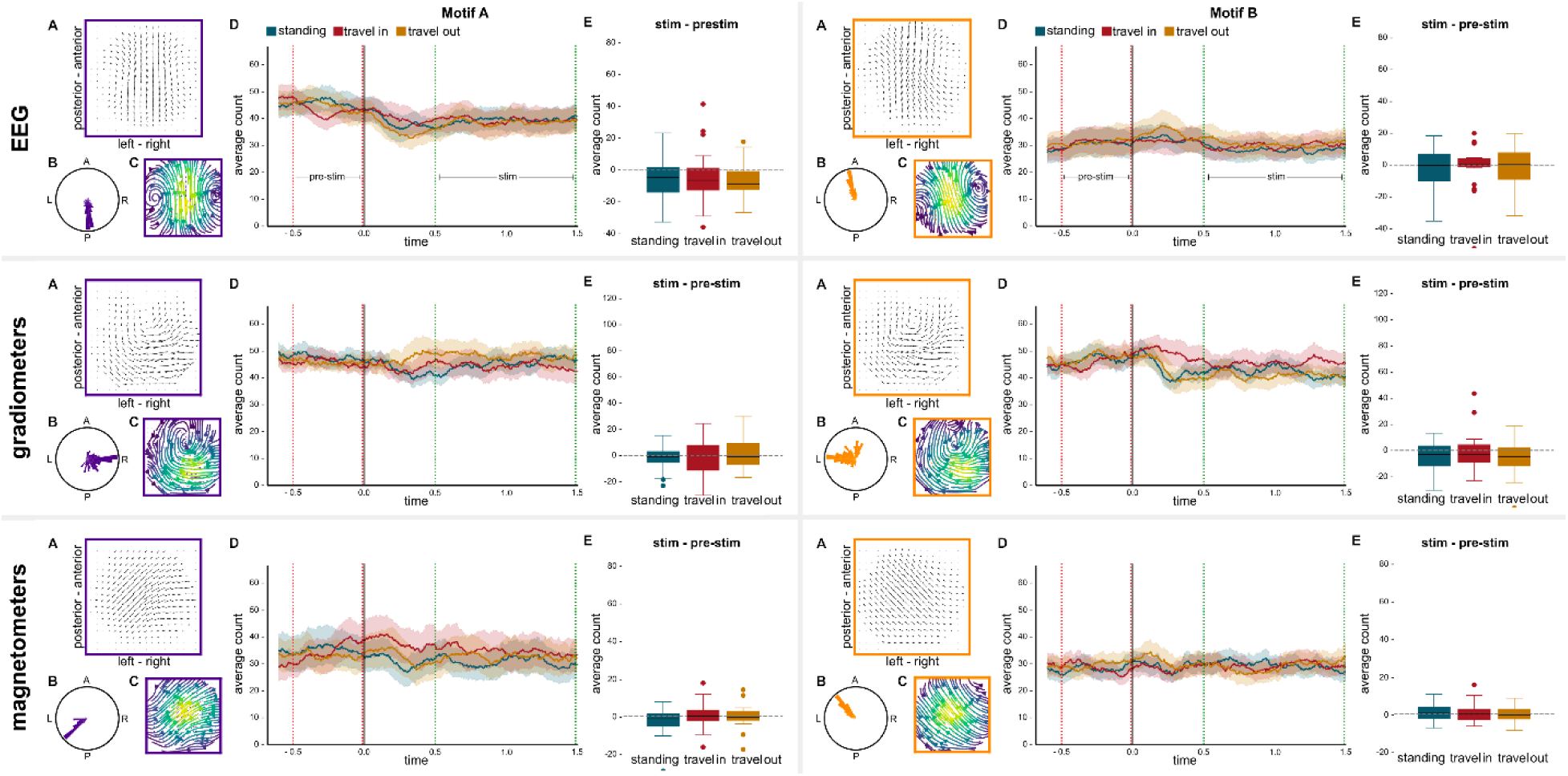
Average counts per condition for the two most frequent motifs (Motif A and Motif B) at participants’ individual alpha frequency.

Overall, stimulus condition was significantly associated with the frequency of occurrence of different motifs during the stimulation period, indicating that motif distribution varied systematically across experimental conditions. This was assessed using a chi-squared test of independence with stimulus condition and counts across trials for each motif in the contingency table, repeated for each timepoint (**Figure 5, upper panel**). To recover granularity in the data, we then considered all six of the most common motifs in a participant-level analysis. A Lasso ridge regression was performed per participant to classify the travel in versus travel out stimulation conditions using the ratio for all motifs. Travel in versus travel out were correctly dissociated in 66 +/-17% of the trials. Note that the classification was above 55% for 12 out of 19 participants, with individual scores up to 98%, highlighting the large variability across participants and sensor types. The specific pattern of motifs explaining conditions similarly varied across participants (**Figure 5, lower panel**).

**Figure 5.**
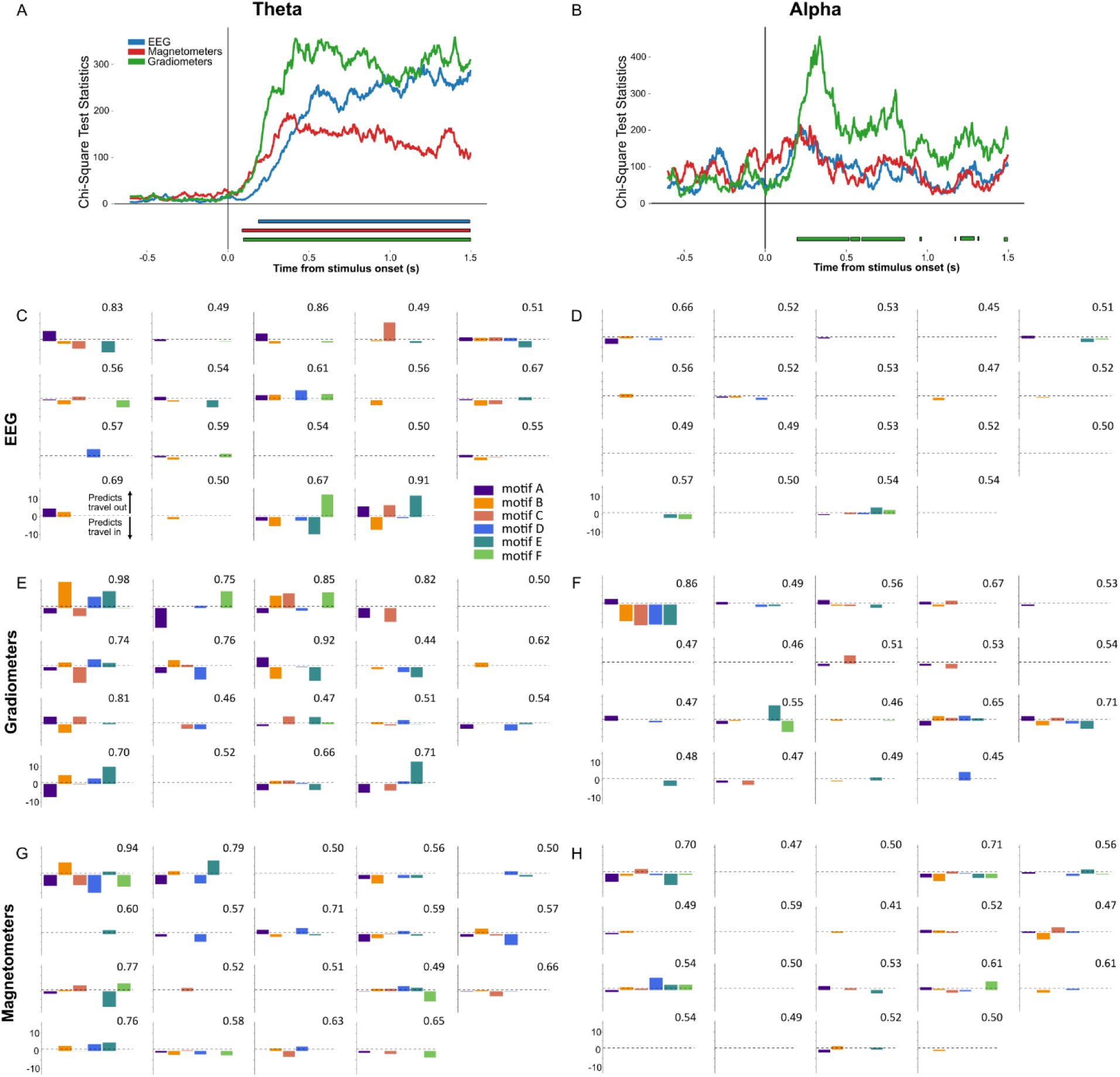
Motifs distributions at 5 Hz discriminate conditions both at the group-level and at the individual level. **A**. Chi-square statistics indicate that the motif counts at 5 Hz differ between conditions during the stimulation period, for all sensor types. **B.** At 10 Hz, only gradiometers show a significant relation of motif counts and stimulation conditions, and primarily during the stimulus onset period. **C, E, G**. The motifs counts were then analyzed for each individual, at 5 Hz and 10 Hz (**D, F, H**), for each sensor type, showing great disparity in terms of distribution. Odds ratio for each motif are represented: odds ratios above 1 indicate that the motif is associated with an increased probability that the trial comes from the travel out condition compared to travel in. The classification accuracy for each participant is indicated in the top right corner of each plot.

### Trial averaging changes apparent wave dynamics

When optical flow was calculated from trial averaged data prior to motif extraction, the motifs that were found strikingly matched those that were derived from individual trials (using the same criteria as for matching within trials, i.e., cosine similarity threshold of 80% in 40% of corresponding grid positions; **Figure 6 A-F**). However, when taking temporal matching into account, i.e., looking at which point in trial-time the motif X occurs, only ∼8% of EEG data, ∼1% of magnetometer and practically none of the gradiometer data matched (for representative example participants’ data and group average see **Figure 6 G-J**).

**Figure 6.**
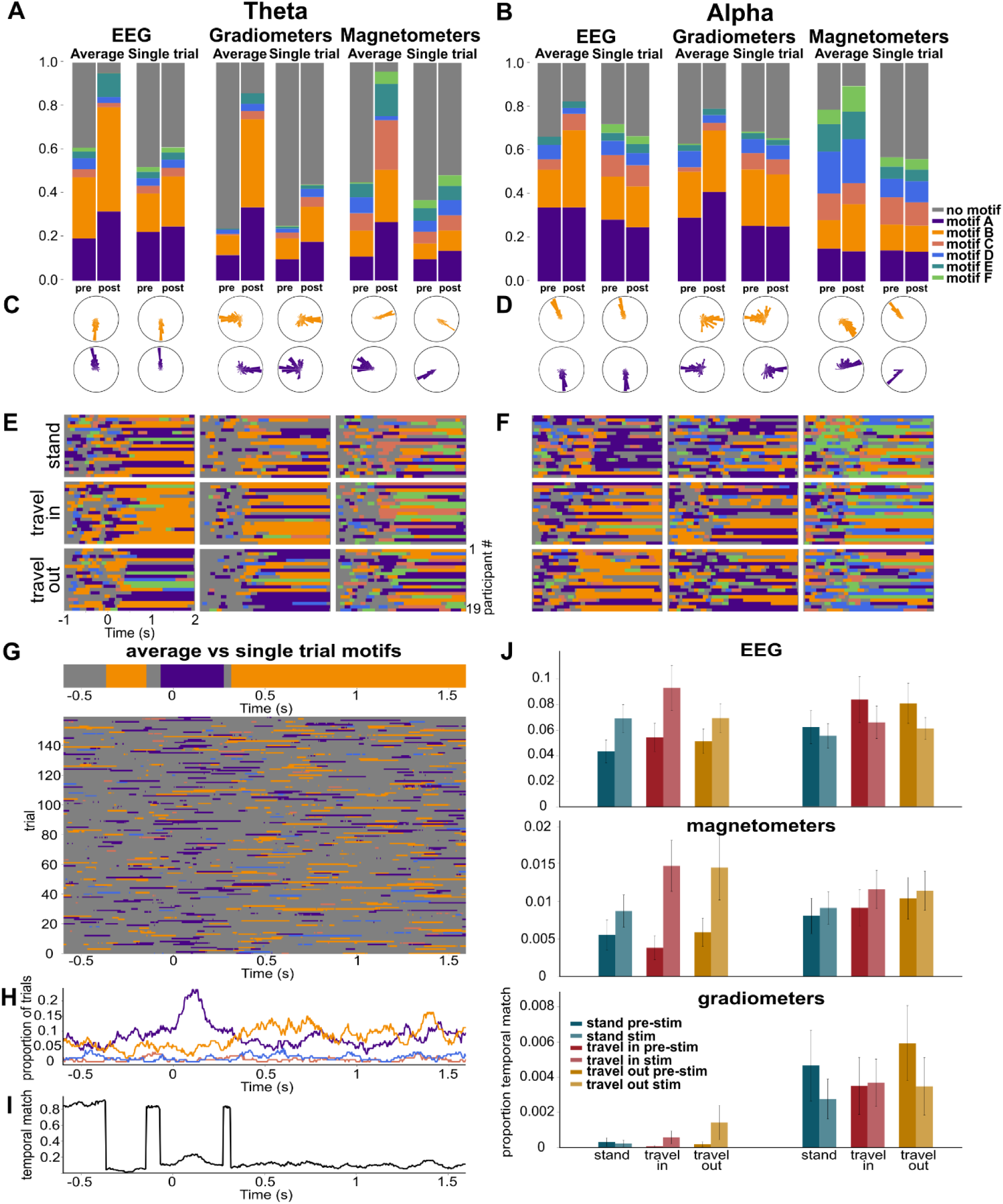
Average versus single trial analysis: A&B. Motif frequency, averaged over all trials and timepoints within the pre-stim and stim intervals, respectively. Motifs were determined from either individually analyzed single trial optical flow maps (‘single trial’) or from optical flow maps based on subject averaged data (‘average’). Analytic signal entering optical flow analysis extracted from theta-band (A) and alpha-band (B) filtered data. **C&D.** Polar plots of motif travel directions derived from theta filtered **(C)** and alpha filtered **(D)** UV maps, respectively. Note that motifs were produced separately for each frequency and averaged vs single trial data, for each sensor type. The motif indices thus follow the order of their occurrence within analysis, but do not indicate similarity across analyses (i.e., motif A in averaged theta optical flow maps is not necessarily the same as motif A in single trial alpha maps). **E&F.** Trial-average motif timeseries, for each participant and condition from theta **(E)** and alpha **(F)** maps. **G.** Comparing motifs from trial averaged data to single trial motifs, example participant. Motifs derived from the trial averaged data (per stimulus condition) of a single participant were used as templates to compare each individual timepoint of each individual trial (from that same participant). Optical flow maps matching any of the average motifs (criteria, see methods), were labelled as that motif. **H.** Proportion of single trials matching average motifs over time, independent of temporal match. **I.** Proportion of individual trials optical flow maps that matched the average motif at the corresponding timepoint. **J.** Bar graphs summarizing the proportion of temporal matches over all participants, for each sensor type and frequency. Note that the scales differ by an order of magnitude between EEG, magnetometers and gradiometers, respectively.

Mesoscopic cortical wave velocities showed minimal differences between participants (mean 0.44m/s, SD 0.09m/s). They were calculated from wave spatial and temporal frequencies, estimated from the stimulus properties in combination with individual fMRI retinotopic maps as 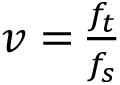 where *f_t_* and *f_s_* are the temporal and spatial frequency of the induced wave in primary visual cortex, respectively.

To estimate wave velocities in the sensor arrays, optical flow vectors of each motif were averaged over all indexed timepoints and projected onto the dominant propagation direction of the corresponding motif. Vector magnitudes, scaled by the geodesic distance along the sensor array and the sampling frequency of the data, were considered as wave velocity estimates.

Given the low variability in the velocities of the visually-induced mesoscopic cortical waves, we did not attempt to correlate them with the estimated velocities of the waves observed in the sensor array. We did, however, compare estimated velocities across motifs as well as stimulation conditions. Due to the uncertainty of optical flow velocity estimates (see methods “Wave velocities”), we caution against strong interpretations of the absolute velocities, although they were well within the range of what is typically reported in EEG studies (**table 1**). Instead, we compared velocity estimates between the most common motifs (motif A and motif B) that corresponded to an EEG sensor array traveling wave from posterior to anterior electrodes and anterior to posterior electrodes, respectively (right to left and left to right, respectively for MEG sensors).

**Table 1.** Wave velocity estimates. Velocities estimated from optical flow vector magnitudes was compared between the two most common motifs (A and B). Travel directions between motifs differ by ∼180°. Reported p-values are uncorrected.

|  | Modality | T-Statistic | P-Value | Motif A |  | Motif B |  |
| --- | --- | --- | --- | --- | --- | --- | --- |
|  |  |  |  | Mean | Std. dev | Mean | Std. dev |
| Theta | EEG | -1.32 | 0.20 | 2.68 | 0.42 | 2.83 | 0.39 |
|  | Gradiometers | 0.81 | 0.43 | 1.21 | 0.19 | 1.15 | 0.27 |
|  | Magnetometers | 0.36 | 0.73 | 2.40 | 0.26 | 2.40 | 0.21 |
|  | Simulations | 0.09 | 0.94 | 3.25 | 0.03 | 3.24 | 0.07 |
| Alpha | EEG | 2.84 | 0.01 | 5.91 | 0.90 | 5.31 | 1.26 |
|  | Gradiometers | -1.53 | 0.14 | 2.14 | 0.52 | 2.35 | 0.31 |
|  | Magnetometers | -1.33 | 0.20 | 4.70 | 0.96 | 5.08 | 0.76 |
|  | Simulations | 1.40 | 0.39 | 2.22 | 0.15 | 1.94 | 0.13 |

Propagation velocities estimated from the optical flow magnitudes did not differ significantly between conditions in any of the comparisons [p>0.05]. Solely the comparison between motif A and motif B in EEG alpha band activity showed a significant uncorrected p-value of p=0.01 (not significant after Bonferroni correction for multiple comparisons; alphaBonf =0.0083).

## Discussion

We induced traveling waves in the retinotopic cortex of 19 research volunteers. From concurrent MEG-EEG recordings, we were able to infer stimulus-wave traveling direction. The directions of the global lags in oscillatory phase at the stimulation frequency across the EEG sensor arrays matched those of the induced traveling wave in primary visual cortex. Travel directions observed in MEG sensors were largely perpendicular to those in EEG sensors, consistent with previous observations (Alexander et al., 2016). We investigated two competing hypotheses about the cortical origins of large-scale traveling waves observed in the extracranial sensor arrays during visual stimulation with a retinotopic traveling wave: (1) projection of mesoscopic cortical waves and (2) sequential region activation along the visual hierarchy (**Figure 1**). The observed condition differences in the corresponding optical flow maps of MEG-EEG phase were consistent with the first hypothesis, i.e., the locally-induced, mesoscopic wave was projected to the global measurement array, there producing an apparent macroscopic wave.

### Limitations and competing explanations

Two considerations challenging this conclusion must be taken into account. First, in single trials, the motifs whose frequency of occurrence differentiated between stimulus conditions during the stimulation period were also present during the pre-stimulus interval, where no stimulus-driven projection could occur. Their frequency (across trials) increased dependent on the stimulation condition (travel in or travel out), compared to the pre-stimulus period. During presentation of the retinotopic standing wave, however, motifs were equally likely to be recorded in anterior-to-posterior as in posterior-to-anterior direction for EEG (left-right to right-left in MEG). Second, considerable differences were observed between participants and across (visually identical) trials, which cannot be readily explained by a straightforward retinotopic mapping of the visual stimulus and its subsequent projection to the sensor array. It is possible, that the criteria for motif membership (85% cosine similarity between corresponding optical flow vectors, equivalent to ∼31.8°) led to the mixing of spontaneously occurring and induced traveling waves.

Under the alternative hypothesis of temporally lagged region activation producing an apparent smooth wave in the sensor array, stimulus condition (travel in vs. travel out vs. standing wave) is not expected to influence the travel direction of the globally observed wave, as the visual hierarchy is preserved across conditions. Instead, the presence versus absence of visual stimulation as a whole is expected to drive wave propagation. Previous studies have explicitly linked traveling wave propagation speeds to axonal conduction delays (Nunez & Srinivasan, 2014). Thus, in the context of sequential activation of regions, wave velocities are expected to differ between motifs (i.e., global wave travel direction) as inter-region temporal delays differ between feedforward and feedback signaling (Markov et al., 2014). Both expectations were contradicted by our data. Stimulus condition significantly influenced global traveling direction and estimated velocities did not differ between motifs with opposite principal directions. Note, however, that alternative accounts of long range traveling waves do not directly rely on inter-region delays (Alexander et al., 2016; Koller et al., 2024; N. Sato, 2022).

In the (non-stimulated) individual participants’ alpha range, we found no condition differences in motif prevalence. The small preference for anterior right to posterior left propagation in gradiometers during the stimulus onset period of the travel in condition (see **Figures 4, middle panel** and **5B**) was outside of the analyzed time periods. More surprisingly, we also found no systematic differences between pre-stimulus and stimulus periods when data were not trial-averaged prior to optical flow estimation. Other studies have instead theoretically predicted (Alamia & VanRullen, 2019) or empirically shown alpha-band traveling wave direction changes with the onset of a visual stimulus (Pang et al., 2020). In both the current, and the previous experiments, the onset of the high contrast inducer stimulus decreased alpha amplitude, thus rendering estimates of oscillatory phase problematic (Cohen, 2014). In our analysis pipeline, all wave motifs were based on individual trial phase maps (where trials with random phase estimates simply resulted in “no motif”). This difference becomes apparent in **Figure 6** where single-trial optical flow analysis of alpha-band filtered data is more likely to result in “no motif” (gray portion of the bar) in the stimulation period, compared to the pre-stimulus period. The opposite is true for the trial-averaged data, where the analytic signal and ergo the optical flow and the derived motifs are biased to high amplitude trials that on average show consistent phase shifts across the sensor array. To rigorously address the ambiguity between alpha phase and amplitude in stimulation versus non-stimulation periods, further careful experimental and modeling work will be required.

### Induced versus spontaneous activity

It is possible that the first harmonic (10 Hz) of the stimulus evoked response to the 5 Hz visual stimulation in our experiment interfered with ongoing oscillatory activity in the alpha band (average individual alpha frequency was mean = 10.06 STD = 0.88). However, if alpha-band responses were contaminated by the first harmonic of the visual stimulation frequency, we would expect similar, albeit weaker, condition effects as seen in the 5 Hz responses. Such effects were not observed in our data.

Previous work in mice has shown that localized, small stimuli produce strong and far reaching (up to full hemispheres) traveling waves (Aggarwal et al., 2022). In contrast, high intensity and widespread stimulation produced weaker traveling waves whose amplitude rapidly decreased with distance from the stimulation site (Sato, Nauhaus, and Carandini 2012). Here, we used high contrast, full-field flicker stimulation to induce retinotopic responses in the shape of a traveling wave. While this method ensures that there is *a* traveling wave in retinotopic cortex, it might have reduced *spontaneously* occurring propagating activity. Thus, in comparing pre-stimulus periods to stimulation periods we might have inadvertently manipulated the likelihood of observing spontaneous traveling waves originating in occipital cortex.

Finally, the strong, frequency-bound, and repetitive full-field visual stimulation applied in our study greatly increased the likelihood of detecting local effects. In contrast, spontaneously occurring, less stereotypical waves more likely lead to the cancellation of local activity, with broader cortical dynamics dominating. Thus, in less constrained experimental conditions, low spatial and temporal frequency activity likely dominates the observed dynamics on the scalp surface (see also Alexander and Dugué 2024).

### Trial averaging

We found that the wave motifs extracted from trial averaged data can only account for ∼8% of timepoints across all individual trials in EEG data (1% in MEG mag, 0.2% in MEG grad). This discrepancy between averaged and single trial data can partially be explained by to the low signal-to-noise ratio inherent to MEG and EEG recordings. Additionally, trial averages are strongly driven by signal amplitude (high amplitude trials weigh higher in the average than low amplitude trials). Conversely, while high amplitude trials dominate the average of the phase estimate, the single-trial derived motifs, are purely based on the unit length orientation of the optical flow vectors and thus nominally indifferent to amplitude variations. Note, however, that phase estimates derived from single-trial Hilbert transforms are notoriously noisy (Cohen 2014) and measures like inter-trial coherence (ITC) are biased towards high-amplitude trials (Aydore et al., 2013) despite being based on trial-by-trial phase estimates. This bias arises because phase estimates for low-amplitude trials tend to be essentially random, whereas those for high-amplitude trials are more reliable. However, in the context of global oscillatory phase shifts, such as those observed in macroscopic traveling waves, we recently demonstrated that between-sensor amplitude correlations are largely independent of the consistency of between-sensor phase lags across epochs (Smulders et al., in prep). Further, intracortical oscillatory traveling waves have been found to be transient (Patten et al., 2012) and temporally variable. Trial averaging might obscure such spatio-temporal dynamics (Alexander et al., 2013) and likely affects faster dynamics disproportionally. We therefore caution against uncritically implementing trial averaging (see also Tlaie et al. 2024).

### Globally observed does not necessarily mean global

Altogether, our results fit well with the hypothesis that the globally observed wave in the sensor array reflects the mesoscopic traveling wave in early visual cortex. However, individual participants’ optical flow patterns showed strong trial-by-trial variability, irrespective of the identical visual stimulation and participants showed strong motif preferences that persisted across stimulation conditions (see individual motif counts, **supplementary figures 1&2**). We conclude that under carefully controlled conditions, non-invasive recordings of global brain activation patterns can reflect localized, mesoscopic traveling waves.

## Acknowledgements

This project has received funding from the European Research Council (ERC) under the European Union’s Horizon 2020 research and innovation programme (grant agreement No 852139 - Laura Dugué). We thank Fren Smulders for his advice on analyses, and Dennis Croonenberg, Yue Kong and David M. Alexander for their useful comments on the manuscript.

---

**Supplementary figure 1:**
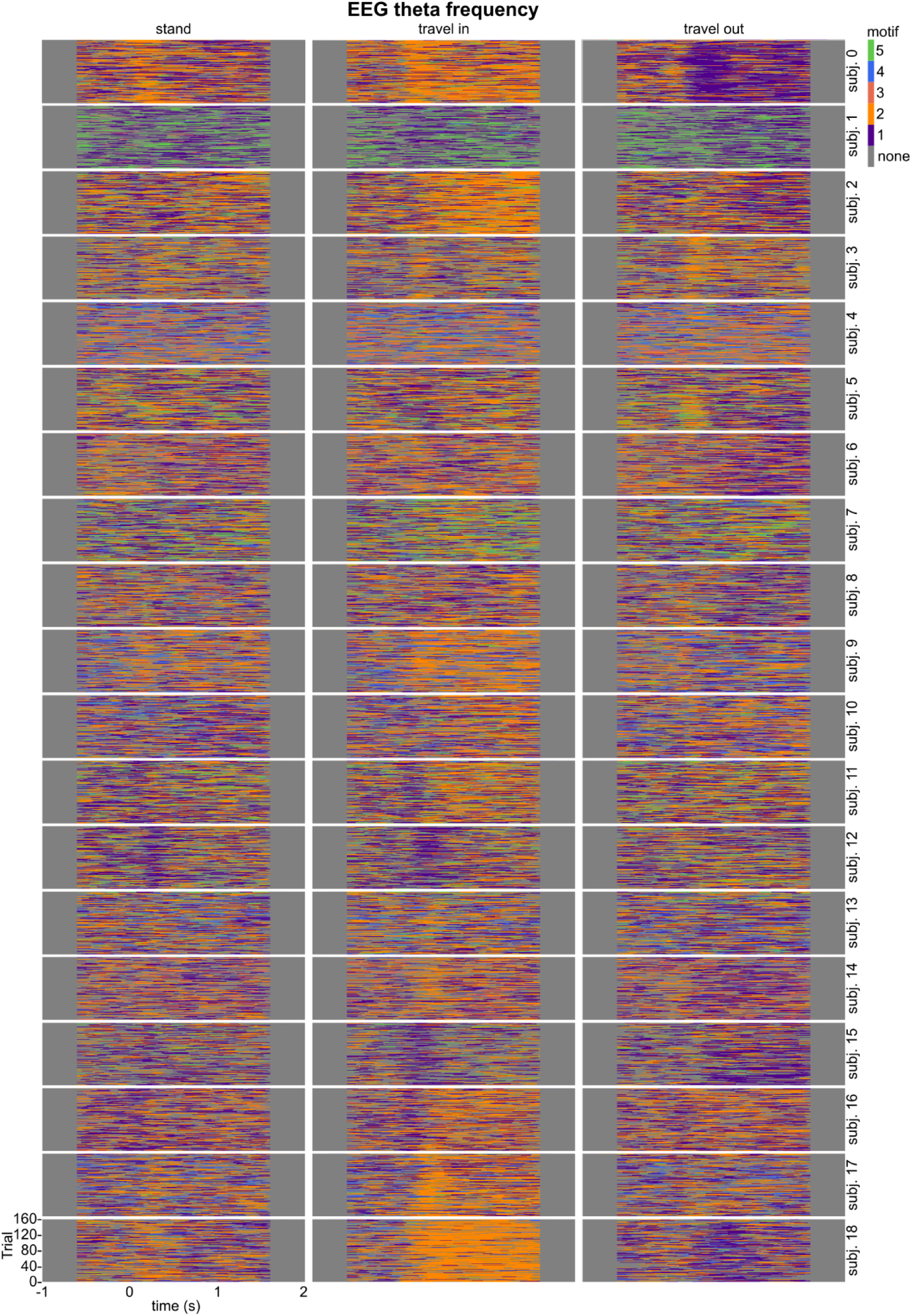
Single participant, single trial motif timeseries EEG theta

**Supplementary figure 2:**
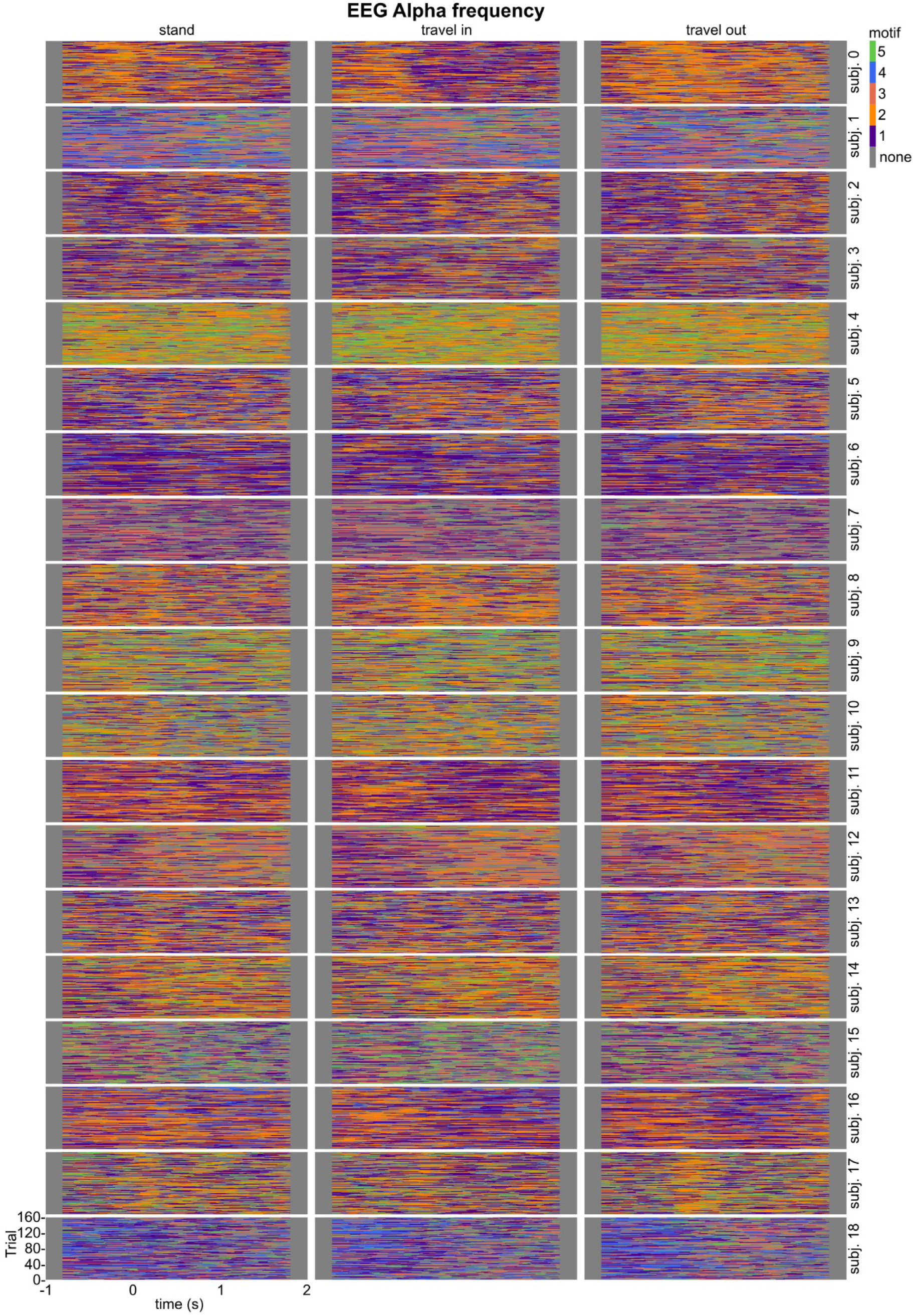
Single participant, single trial motif timeseries EEG alpha

## References

1. Aggarwal, A., Brennan, C., Luo, J., Chung, H., Contreras, D., Kelz, M. B., & Proekt, A. (2022). Visual evoked feedforward–feedback traveling waves organize neural activity across the cortical hierarchy in mice. Nature Communications, 13(1), 4754.

2. Alamia, A., Terral, L., D’ambra, M. R., & VanRullen, R. (2023). Distinct roles of forward and backward alpha-band waves in spatial visual attention. Elife, 12, e85035.

3. Alamia, A., & VanRullen, R. (2019). Alpha oscillations and traveling waves: Signatures of predictive coding? PLoS Biology, 17(10), e3000487.

4. Alexander, D. M., & Dugué, L. (2024). The dominance of global phase dynamics in human cortex, from delta to gamma. 10.7554/elife.100674.1

5. Alexander, D. M., Jurica, P., Trengove, C., Nikolaev, A. R., Gepshtein, S., Zvyagintsev, M., Mathiak, K., Schulze-Bonhage, A., Ruescher, J., & Ball, T. (2013). Traveling waves and trial averaging: The nature of single-trial and averaged brain responses in large-scale cortical signals. Neuroimage, 73, 95–112.

6. Alexander, D. M., Nikolaev, A. R., Jurica, P., Zvyagintsev, M., Mathiak, K., & van Leeuwen, C. (2016). Global neuromagnetic cortical fields have non-zero velocity. PLoS One, 11(3), e0148413.

7. Alexander, D. M., Trengove, C., Wright, J. J., Boord, P. R., & Gordon, E. (2006). Measurement of phase gradients in the EEG. Journal of Neuroscience Methods, 156(1–2), 111–128.

8. Aydore, S., Pantazis, D., & Leahy, R. M. (2013). A note on the phase locking value and its properties. Neuroimage, 74, 231–244.

9. Barron, J. L., Fleet, D. J., & Beauchemin, S. S. (1994). Performance of optical flow techniques. International Journal of Computer Vision, 12, 43–77.

10. Borg, I., & Groenen, P. J. (2005). Modern multidimensional scaling: Theory and applications. Springer Science & Business Media.

11. Broderick, W. F., Simoncelli, E. P., & Winawer, J. (2022). Mapping spatial frequency preferences across human primary visual cortex. Journal of Vision, 22(4), 3–3.

12. Bullier, J. (2001). Integrated model of visual processing. Brain Research Reviews, 36(2), 96–107.

13. Busch, N. A., Dubois, J., & VanRullen, R. (2009). The phase of ongoing EEG oscillations predicts visual perception. Journal of Neuroscience, 29(24), 7869–7876.

14. Cohen, M. X. (2014). Analyzing neural time series data: Theory and practice. MIT press.

15. Donoghue, T., Haller, M., Peterson, E. J., Varma, P., Sebastian, P., Gao, R., Noto, T., Lara, A. H., Wallis, J. D., & Knight, R. T. (2020). Parameterizing neural power spectra into periodic and aperiodic components. Nature Neuroscience, 23(12), 1655–1665.

16. Dugué, L., Marque, P., & VanRullen, R. (2011). The phase of ongoing oscillations mediates the causal relation between brain excitation and visual perception. Journal of Neuroscience, 31(33), 11889–11893.

17. Dumoulin, S. O., & Wandell, B. A. (2008). Population receptive field estimates in human visual cortex. Neuroimage, 39(2), 647–660.

18. Esteban, O., Markiewicz, C. J., Blair, R. W., Moodie, C. A., Isik, A. I., Erramuzpe, A., Kent, J. D., Goncalves, M., DuPre, E., & Snyder, M. (2019). fMRIPrep: A robust preprocessing pipeline for functional MRI. Nature Methods, 16(1), 111–116.

19. Fakche, C., & Dugué, L. (2024). Perceptual cycles travel across retinotopic space. Journal of Cognitive Neuroscience, 36(1), 200–216.

20. Fakche, C., Galas, L., Petras, K., & Dugué, L. (2024). Alpha traveling waves index spatial attention. Vision Sciences Society 24th annual Meeting.

21. Fakche, C., VanRullen, R., Marque, P., & Dugué, L. (2022). α phase-amplitude tradeoffs predict visual perception. ENeuro, 9(1).

22. Felleman, D. J., & Van, D. C. E. (1991). Distributed hierarchical processing in the primate cerebral cortex. Cerebral Cortex (New York, NY: 1991), 1(1), 1–47.

23. Friedman, J., Hastie, T., & Tibshirani, R. (2010). Regularization paths for generalized linear models via coordinate descent. Journal of Statistical Software, 33(1), 1.

24. Grabot, L., Merholz, G., Winawer, J., Heeger, D. J., & Dugué, L. (2024). Traveling Waves in the Human Visual Cortex: A MEG-EEG Model-Based Approach. bioRxiv, 2024.10.09.617389. 10.1101/2024.10.09.617389

25. Gramfort, A., Luessi, M., Larson, E., Engemann, D. A., Strohmeier, D., Brodbeck, C., Parkkonen, L., & Hämäläinen, M. S. (2014). MNE software for processing MEG and EEG data. Neuroimage, 86, 446–460.

26. Gutzen, R., De Bonis, G., De Luca, C., Pastorelli, E., Capone, C., Mascaro, A. L. A., Resta, F., Manasanch, A., Pavone, F. S., & Sanchez-Vives, M. V. (2024). A modular and adaptable analysis pipeline to compare slow cerebral rhythms across heterogeneous datasets. Cell Reports Methods, 4(1).

27. Hillebrand, A., & Barnes, G. R. (2005). Beamformer analysis of MEG data. International Review of Neurobiology, 68, 149–171.

28. Hillebrand, A., & Barnes, G. R. (2011). Practical constraints on estimation of source extent with MEG beamformers. Neuroimage, 54(4), 2732–2740.

29. Hindriks, R., van Putten, M. J., & Deco, G. (2014). Intra-cortical propagation of EEG alpha oscillations. Neuroimage, 103, 444–453.

30. Horn, B. K., & Schunck, B. G. (1981). Determining optical flow. Artificial Intelligence, 17(1–3), 185–203.

31. Horton, J. C., & Hoyt, W. F. (1991). The representation of the visual field in human striate cortex: A revision of the classic Holmes map. Archives of Ophthalmology, 109(6), 816– 824.

32. Kay, K. N., Winawer, J., Mezer, A., & Wandell, B. A. (2013). Compressive spatial summation in human visual cortex. Journal of Neurophysiology, 110(2), 481–494.

33. Kleiner, M., Brainard, D., & Pelli, D. (2007). What’s new in Psychtoolbox-3?

34. Koller, D. P., Schirner, M., & Ritter, P. (2024). Human connectome topology directs cortical traveling waves and shapes frequency gradients. Nature Communications, 15(1), 3570.

35. Kriegeskorte, N., Mur, M., Ruff, D. A., Kiani, R., Bodurka, J., Esteky, H., Tanaka, K., & Bandettini, P. A. (2008). Matching categorical object representations in inferior temporal cortex of man and monkey. Neuron, 60(6), 1126–1141.

36. Lin, F.-H., Witzel, T., Zeffiro, T. A., & Belliveau, J. W. (2008). Linear constraint minimum variance beamformer functional magnetic resonance inverse imaging. Neuroimage, 43(2), 297–311.

37. Markov, N. T., Vezoli, J., Chameau, P., Falchier, A., Quilodran, R., Huissoud, C., Lamy, C., Misery, P., Giroud, P., & Ullman, S. (2014). Anatomy of hierarchy: Feedforward and feedback pathways in macaque visual cortex. Journal of Comparative Neurology, 522(1), 225–259.

38. Mathewson, K. E., Gratton, G., Fabiani, M., Beck, D. M., & Ro, T. (2009). To see or not to see: Prestimulus α phase predicts visual awareness. Journal of Neuroscience, 29(9), 2725– 2732.

39. Mouraux, A., & Iannetti, G. D. (2008). Across-trial averaging of event-related EEG responses and beyond. Magnetic Resonance Imaging, 26(7), 1041–1054.

40. Muller, L., Chavane, F., Reynolds, J., & Sejnowski, T. J. (2018). Cortical travelling waves: Mechanisms and computational principles. Nature Reviews Neuroscience, 19(5), 255– 268.

41. Nunez, P. L., & Srinivasan, R. (2014). Neocortical dynamics due to axon propagation delays in cortico-cortical fibers: EEG traveling and standing waves with implications for top-down influences on local networks and white matter disease. Brain Research, 1542, 138–166.

42. Orczyk, J. J., & Kajikawa, Y. (2022). Magnifying traveling waves on the scalp. Brain Topography, 35(1), 162–168.

43. Orsher, Y., Rom, A., Perel, R., Lahini, Y., Blinder, P., & Shein-Idelson, M. (2023). Travelling waves or sequentially activated discrete modules: Mapping the granularity of cortical propagation. bioRxiv, 2023–08.

44. Pang, Z., Alamia, A., & VanRullen, R. (2020). Turning the stimulus on and off changes the direction of α traveling waves. Eneuro, 7(6).

45. Patten, T. M., Rennie, C. J., Robinson, P. A., & Gong, P. (2012). Human cortical traveling waves: Dynamical properties and correlations with responses. PLoS One, 7(6), e38392.

46. Sato, N. (2022). Cortical traveling waves reflect state-dependent hierarchical sequencing of local regions in the human connectome network. Scientific Reports, 12(1), 334.

47. Sato, T. K., Nauhaus, I., & Carandini, M. (2012). Traveling waves in visual cortex. Neuron, 75(2), 218–229.

48. Smulders, F.T.Y., Petras, K., & ten Oever, S., Dissociating components of EEG coherence. (in preparation)

49. Sokoliuk, R., & VanRullen, R. (2016). Global and local oscillatory entrainment of visual behavior across retinotopic space. Scientific Reports, 6(1), 25132.

50. Strasburger, H., Rentschler, I., & Jüttner, M. (2011). Peripheral vision and pattern recognition: A review. Journal of Vision, 11(5), 13–13.

51. Taulu, S., & Kajola, M. (2005). Presentation of electromagnetic multichannel data: The signal space separation method. Journal of Applied Physics, 97(12).

52. Tlaie, A., Shapcott, K., van der Plas, T. L., Rowland, J., Lees, R., Keeling, J., Packer, A., Tiesinga, P., Schölvinck, M. L., & Havenith, M. N. (2024). What does the mean mean? A simple test for neuroscience. PLOS Computational Biology, 20(4), e1012000.

53. Townsend, R. G., & Gong, P. (2018). Detection and analysis of spatiotemporal patterns in brain activity. PLoS Computational Biology, 14(12), e1006643.

54. van den Broek, S. P., Reinders, F., Donderwinkel, M., & Peters, M. J. (1998). Volume conduction effects in EEG and MEG. Electroencephalography and Clinical Neurophysiology, 106(6), 522–534.

55. Van Essen, D. C., & Maunsell, J. H. R. (1983). Hierarchical organization and functional streams in the visual cortex. Trends in Neurosciences, 6, 370–375.

56. Vanpoucke, F. J., Boermans, P.-P. B., & Frijns, J. H. (2011). Assessing the placement of a cochlear electrode array by multidimensional scaling. IEEE Transactions on Biomedical Engineering, 59(2), 307–310.

57. Widmann, A., Schröger, E., & Maess, B. (2015). Digital filter design for electrophysiological data–a practical approach. Journal of Neuroscience Methods, 250, 34–46.

58. Wolters, C. H., Anwander, A., Maess, B., Macleod, R. S., & Friederici, A. D. (2004). The influence of volume conduction effects on the EEG/MEG reconstruction of the sources of the Early Left Anterior Negativity. The 26th Annual International Conference of the IEEE Engineering in Medicine and Biology Society, 2, 3569–3572.

59. Zhang, H., Watrous, A. J., Patel, A., & Jacobs, J. (2018). Theta and alpha oscillations are traveling waves in the human neocortex. Neuron, 98(6), 1269–1281.

60. Zhigalov, A., & Jensen, O. (2023). Travelling waves observed in MEG data can be explained by two discrete sources. NeuroImage, 272, 120047.

